# Hepatic Leukemia Factor supports the propagation of leukemia and hematopoietic stem cell function during stress-induced regeneration

**DOI:** 10.1101/2021.03.05.434034

**Authors:** Aurélie Baudet, Karolina Komorowska, Simon Hultmark, Marion Chapellier, Kenichi Miharada, Jonas Larsson, David Bryder, Marcus Järås, Gunnar Juliusson, Mattias Magnusson

## Abstract

The processes regulating hematopoietic stem cells (HSC) during aging are not fully understood^1^, but it is clear that the incidence of hematological malignancies increases with age, highlighting the importance of unravelling the cellular and molecular networks involved. Recently, we identified Hepatic Leukemia Factor (HLF) as an essential transcription factor in maintaining the HSC pool during regeneration^2^ and showed that failure to downregulate HLF leads to disrupted differentiation^3^.

Here, we found that HLF is dispensable for hematopoiesis during systemic aging, but needed during stress-induced hematopoietic recovery of aged HSC after transplantation. Additionally, HLF was dispensable for leukemic initiation but required for disease propagation. Taken together, our findings demonstrate the existence of a HLF-dependent mechanism that uncouples stress-induced regeneration from hematopoietic homeostasis during aging, that can be used by malignant cells to gain stem cell properties to propagate the disease.

**Key points:**

- HLF is dispensable for HSC function and hematopoietic homeostasis during physiological aging, but crucial during stress induced regeneration.
- HLF supports the propagation of leukemia-initiating cells

## Methods

### Mice

The generation of KO mice was previously described^4^, and mice were backcrossed to achieve pure C57BL/6 background. Animals were housed in ventilated racks, given autoclaved food and water, and maintained in accordance with Swedish Animal Welfare organisation guidelines, at the Biomedical Center animal facilities in Lund. All animal experiments were approved by local ethical committees (permit M94-15).

### Competitive transplantation assay

For transplantation, 2 x10^5^ unfractionated cells from BM from 18-month-old mice (CD45.2) were mixed in a 1:1 ratio with 2×10^5^ unfractionated BM competitor cells (CD45.1). For secondary transplant, a femur per donor was split into 2 recipients. Grafts were intravenously injected into lethally irradiated (900 cGy) recipients (CD45.1/CD45.2).

### Generation of MLL/AF9 leukemia

MLL/AF9 leukemia was generated as described in^5^ from either WT or KO cKit^+^ cells. HLF expression was established by semi-quantitative PCR as described in^2^.

### Peripheral blood and bone marrow preparation

Peripheral blood (PB) was collected from the tail vein. Blood parameters were analyzed using SysmexXE-5000 (Sysmex Europe GmbH). Before staining, erythrocytes were lysed with NH4Cl (StemCell Technologies). Bone marrow cells were isolated by crushing the femur and tibia in PBS containing 2% FCS (GIBCO), and passed through a 70 μm filter.

### Flow cytometry analysis

All experiments were performed using FACS Canto, FACS LSR II cytometers, FACS Aria IIu or FACS Aria III cell sorters (Becton Dickinson) and analyzed by FlowJo software (Tree Star). Antibodies used in analyses are listed in Table S1.

### Statistical analyses

Statistical analysis was performed with GraphPad Prism software. Error bars represent SEM.

## Results and discussion

HSC support blood production throughout life. Still, their regenerative capacity declines with age^6^. Since HLF has previously been found to be crucial for maintaining HSC function in adult mice (12-16 weeks) during regeneration^2^, we here investigated the role of HLF in HSC during systemic aging and in acute myeloid leukemia, the latter representing an age-associated disease. Knockout (KO) of HLF in mice did not appear to influence their overall lifespan, as survival at 18 month of age was over 90% for both KO mice and WT littermate controls (n>10). Aged KO mice displayed normal hematopoietic parameters except for a reduction in blood platelets (Sup Fig. 1A), which is in line with our previous findings in younger adult mice^2^. Furthermore, the lineage distribution in peripheral blood (PB) and bone marrow (BM) of aged KO mice revealed an increased myeloid output (Fig1A, 1B) in comparison to younger adult KO mice, demonstrating a normal aging phenotype^7, 8^. The expression levels of HLF with age remained conserved in hematopoietic stem and progenitor cells in comparison to younger adult mice (Fig1C and ^2^), establishing that the phenotype was not due to aging-related systemic changes in gene expression. We next investigated the immunophenotypic frequency and function of aged KO HSC. No significant difference was observed in the frequency of immature hematopoietic progenitor (LSK, Lin^-^ Sca^+^ c-Kit^+^) cells or phenotypic HSC in the aged KO mice compared to WT controls (Fig1D). Despite an increase in the cycling pool of HSC observed in younger adult mice, all KO-derived aged HSC were found to be quiescent (Fig.1E) at levels similar to WT mice^9^, further underlining the redundancy of HLF during systemic aging. Collectively, these data demonstrated that HLF is dispensable for native hematopoiesis during aging. However, in serial competitive transplantations, the KO cells displayed a reduction in reconstitution capacity in primary recipients that became more pronounced in secondary recipients (Fig1F), while the lineage distribution remained normal (Fig.1G). This highlights a central role for HLF in uncoupling stress-induced hematopoiesis from the continuous hematopoietic regeneration during aging. This suggests that HLF may control a sub-program of the HSC that governs symmetric self-renewal during the HSC expansion needed after transplantation or insults, while dispensable for steady state hematopoiesis. Alternatively, the loss of quiescence observed in KO HSC in adult mice may exhaust the capacity of the HSC to respond to external insult to the point that they can only accommodate the proliferative requirement of steady-state hematopoiesis.

**Figure 1:**
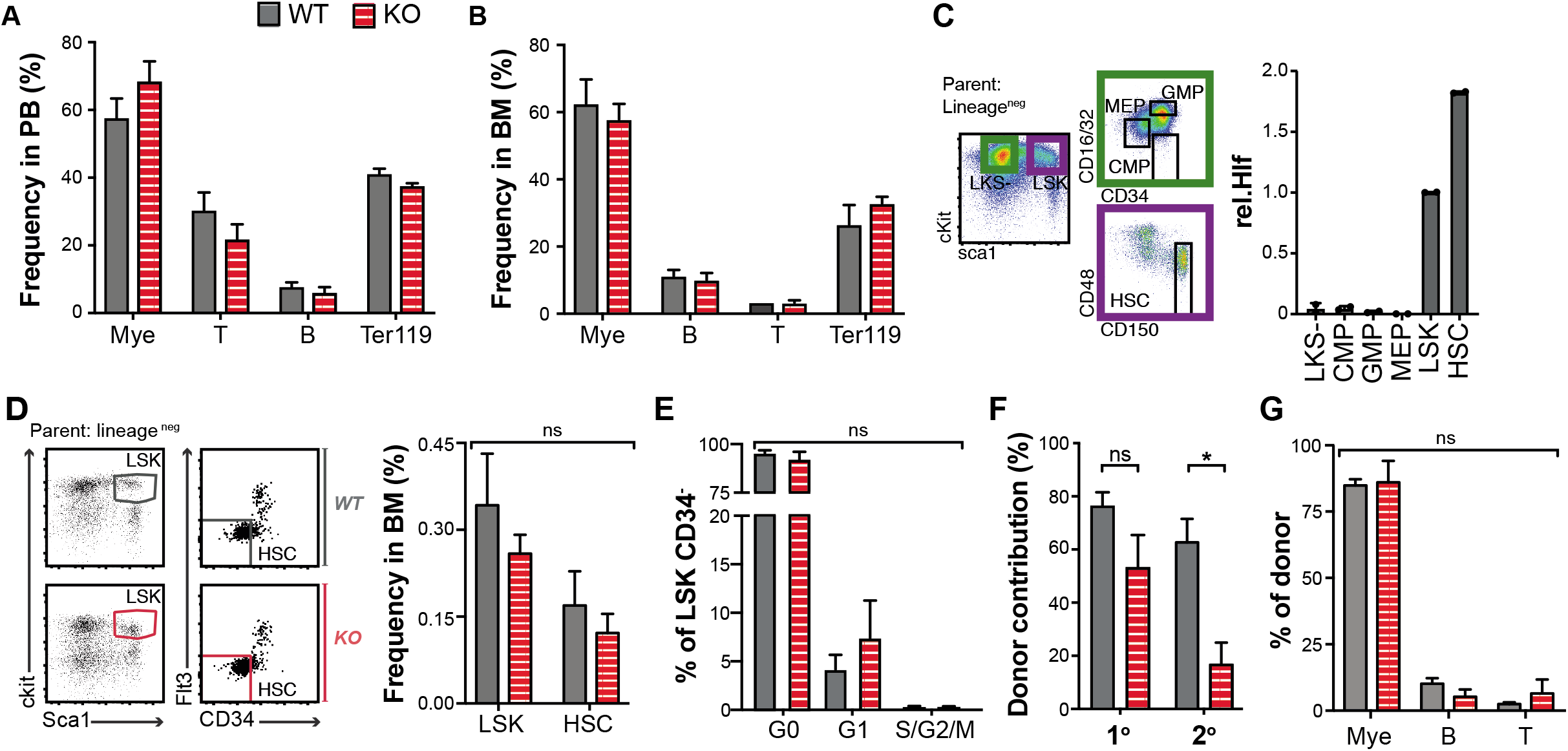
HLF uncouples stress-regeneration from steady-state production during systemic hematopoietic aging. A/ and B/ Myeloid (Mye, Gr1^+^Mac1^+^), B- (B220^+^), T- (CD3^+^) cells, and erythrocytes (Ter119^+^) percentage in (A) the PB and, (B) the BM of WT or KO 18-month old mice. C/ Relative expression of HLF/HPRT in hematopoietic stem and progenitor compartments sorted from the BM of 18 month-old mice. D/ Hematopoietic progenitors (LSK, Lineage^-^cKit^+^sca1^+^) and stem cells (HSC, LSK-CD34^-^flt3^-^) percentage in the bone marrow of 18 month-old WT (n=9) and KO (n=10) mice. E/ Ki67 staining of HSC in the BM of WT and KO 18 month-old mice. F/ Donor contribution in BM after 20 and 16 weeks respectively in primary (1°) and secondary (2°) recipients of transplantation of BM from either WT or KO 18-month mice (3 donors per genotype). G/ Lineage distribution in the BM of engrafted 2° recipients in E. Legend; PB: periphearl blood, BM: bone marrow, WT: wild type, KO: double HLF knock-out, ns: non-significant, * p<0,05. All data acquired with FACS. FACS plots show representative profile. Unpaired t-test was used to evaluate

Next, as many studies have shown a role for HLF in AML^10, 11, 12^, an aging-related disease, we hypothesized that HLF may potentially sustain the leukemic stem cells during the propagation of disease, in addition to its role in maintaining HSC during regeneration. To address this, we retrovirally expressed *MLL/AF9 (MA9)* in bone marrow-derived c-Kit^+^ cells from KO and WT mice, and transplanted them into syngeneic recipient mice (Fig.2A)^5^. Although no difference was observed in the latency or overall survival between the primary recipient groups (Fig.2B and 2C), secondary recipient mice receiving HLF KO *MA9* cells lived significantly longer than mice receiving WT *MA9* cells (Fig.2D). This demonstrates that HLF is dispensable for the initiation of *MA9*-induced AML but has a role for disease propagation. Given that the tumorinitiating cells in this disease model initially lack expression of HLF^13^, we hypothesized that upon propagation of the disease, HLF expression becomes up-regulated to allow the disease cell to rapidly expand. To test this, we performed semi-quantitative PCR on the WT leukemic cells, showing a clear increase of HLF expression in AML cells from the secondary recipient mice compared to those from primary recipients (Fig.2E). Altogether, these data propose that HLF expression can be “rewired”, potentially as a strategy for the leukemia to acquire stem cell properties and subsequently promote propagation of the disease. As such, HLF could be considered a possible therapeutic target to inhibit disease progression. Overall, our results highlight an HLF-dependent mechanism that maintains leukemic stem cells and HSC function during stress-induced regeneration which appears uncoupled from native hematopoiesis during aging.

**Figure 2:**
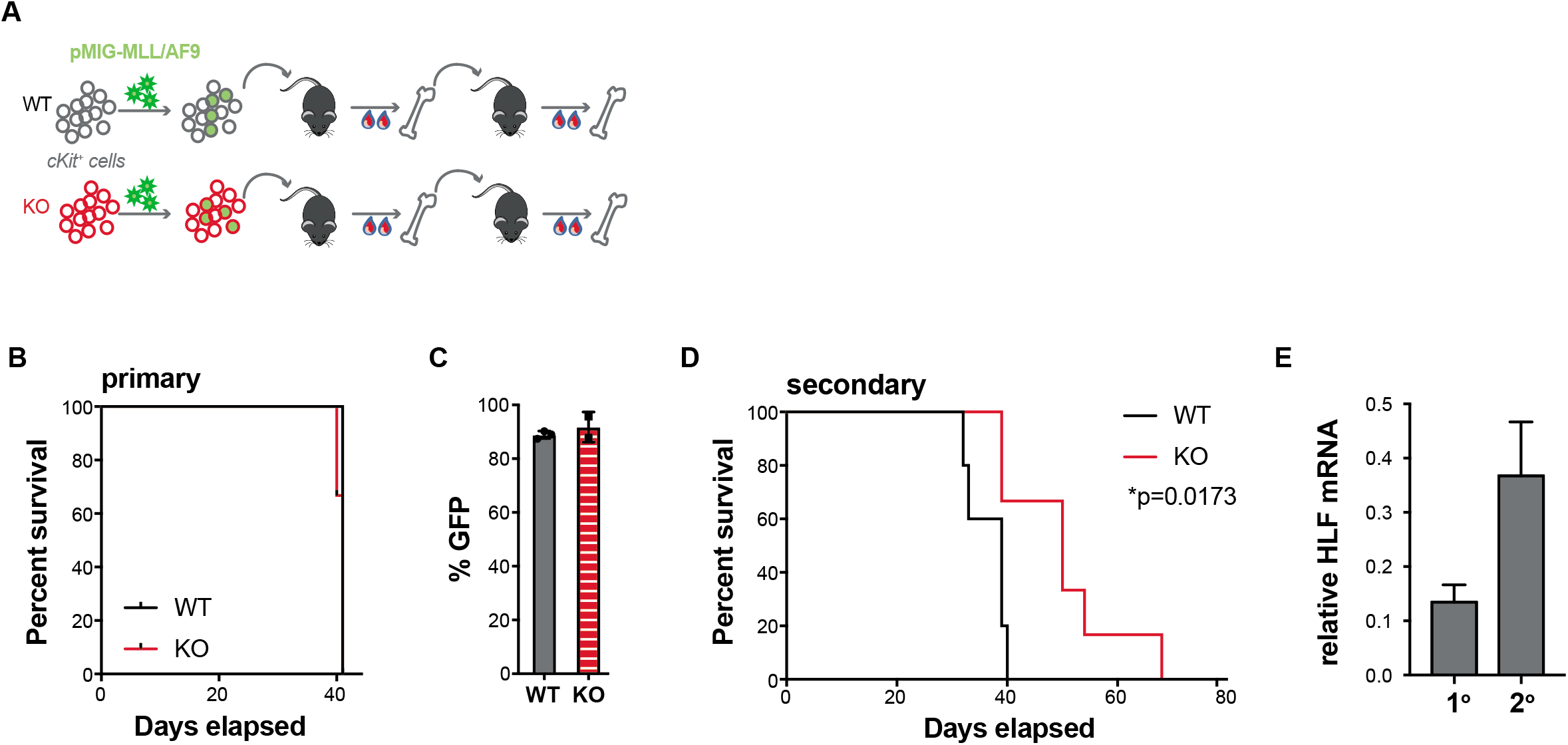
HLF promotes MLL/AF9-dependent leukemogenesis. A/ Study design. B/ Survival curve of primary recipients of either WT or KO MLL/AF9-transduced cKit^+^ cells. C/ Percentage of GFP^+^ cells was established by FACS in the BM of surviving animals at day 40 (see B). D/ Survival curve of secondary recipients of either WT or KO MLL/AF9 leukemia from animals in B. E/ The expression of HLF, normalized to HPRT, was quantified in the BM of recipients of WT leukemia. Legend; BM: bone marrow, WT: wild type, KO: double HLF knock-out, 1°/2°: primary/secondary recipients. Animals were sacrificed at humane end point. Data (C and E) are shown as mean plus SEM. Mantel-Cox test was use in D.

## Acknowledgements

The authors would like to thank support staff at the Lund Stem Cell Center, Lund University as well as their funding sources (Swedish Childhood Cancer Foundation, Swedish Cancer Foundation, and Swedish Research Council).

## Authorship Contributions

AB, KK, and SH designed, and performed experiments. MC and MJ provided MLL/AF9 retrovirus. AB, and MM wrote the manuscript. All authors contributed to study design and manuscript correction.

## Disclosure of Conflicts of Interest

No conflict to disclose.

---

**Supplemental figure 1:**
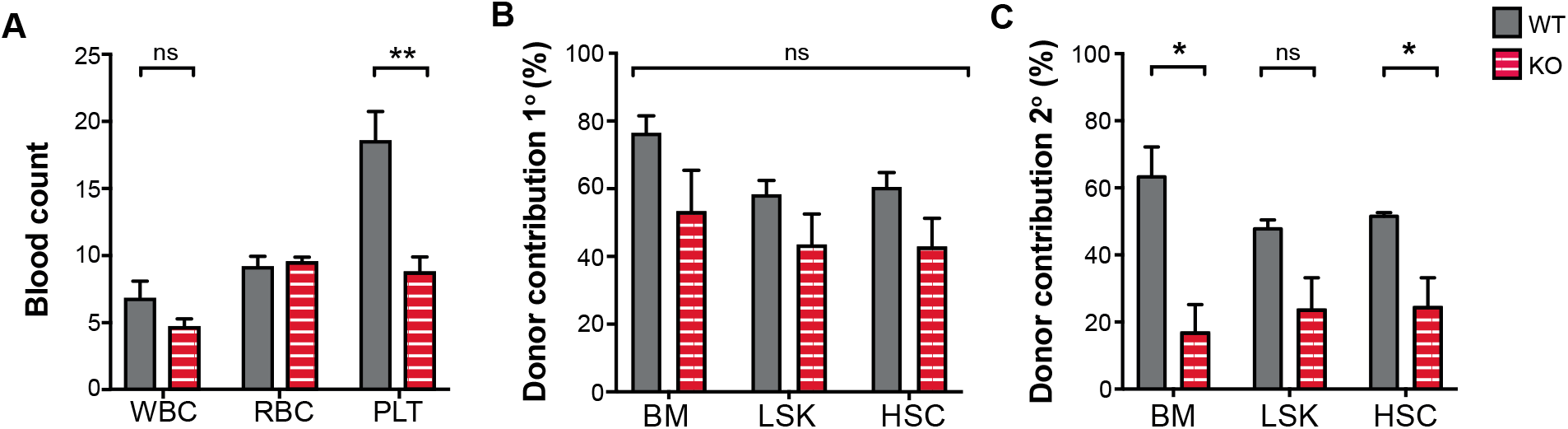
Effect of HLF KO on the hematopoiesis of aged animals. A/ Sysmex counts in the PB of 18 month-old mice. B and C/ Donor contribution was quantified in specified compartments after 20 and 16 weeks respectively in primary (B) and secondary (C) recipients of transplantation of BM from either 18-month-old WT or KO mice (3 donors per genotype). Legend; PB: Peripheral blood, BM: bone marrow, WT: wild type, KO: HLF knock-out, WBC: white blood cells x10^9^, RBC: red blood cells x10^12^, PLT: platelets x10^11^, ns: non-significant, ** p<0,01, * p<0,05. All data acquired with FACS. Unpaired t-test was used to evaluate statistical significance.

## Notes

### Competing Interest Statement

The authors have declared no competing interest.

